# A microvolume method for measuring catalase activity

**DOI:** 10.1101/2025.11.07.687299

**Authors:** Beatriz Corrêa da Rocha, Marcos Trindade Da Rosa, João Batista Teixeira Da Rocha, Elgion Lucio Silva Loreto

**Affiliations:** Center for Natural and Exact Sciences, Doctoral Program Toxicological Biochemistry, Universidade Federal de Santa Maria (UFSM), CEP, Santa Maria, RS, Brazil; Center for Natural and Exact Sciences, Doctoral Program in Animal Biodiversity, Universidade Federal de Santa Maria (UFSM), Santa Maria, RS, Brazil; Department of Biochemistry and Molecular Biology, Center for Natural and Exact Sciences, Federal University of Santa Maria, Camobi, Santa Maria, Av Roraima, 1000, 97105-900, Brazil

**Keywords:** Catalase activity protocol, NanoDrop, microvolume, Oxidative stress, nanospectrophotometer

## Abstract

We have developed and validated an innovative protocol for analyzing catalase activity in microvolumes using the NanoDrop spectrophotometer. This method offers a solution to the challenge of working with limited biological samples and provides an efficient alternative to conventional protocols that require larger sample volumes. Unlike typical microplate assays that aim to increase throughput or reduce costs, our protocol was developed specifically for scenarios where biological material is scarce, such as studies with small organisms like *Panagrellus redivivus* and *Caenorhabditis elegan*s, as well as with certain tissues of *Drosophila* and other small organisms. A key advantage of the method described here is the ability to accurately measure catalase activity with as little as 2 μL of sample, making it ideal for studies where sample availability is extremely limited. The results show that the protocol effectively assesses catalase efficiency and reflects the physiological and metabolic properties of the tissues studied. Inhibitors and denaturants were used to ensure specificity of catalase measurements and the method was optimized for minimal reagent consumption. This approach greatly expands the research possibilities on enzymatic activity in reduced biological models, especially in contexts where small samples are critical, such as limited tissue collections or small organisms

**Graphical abstract:** 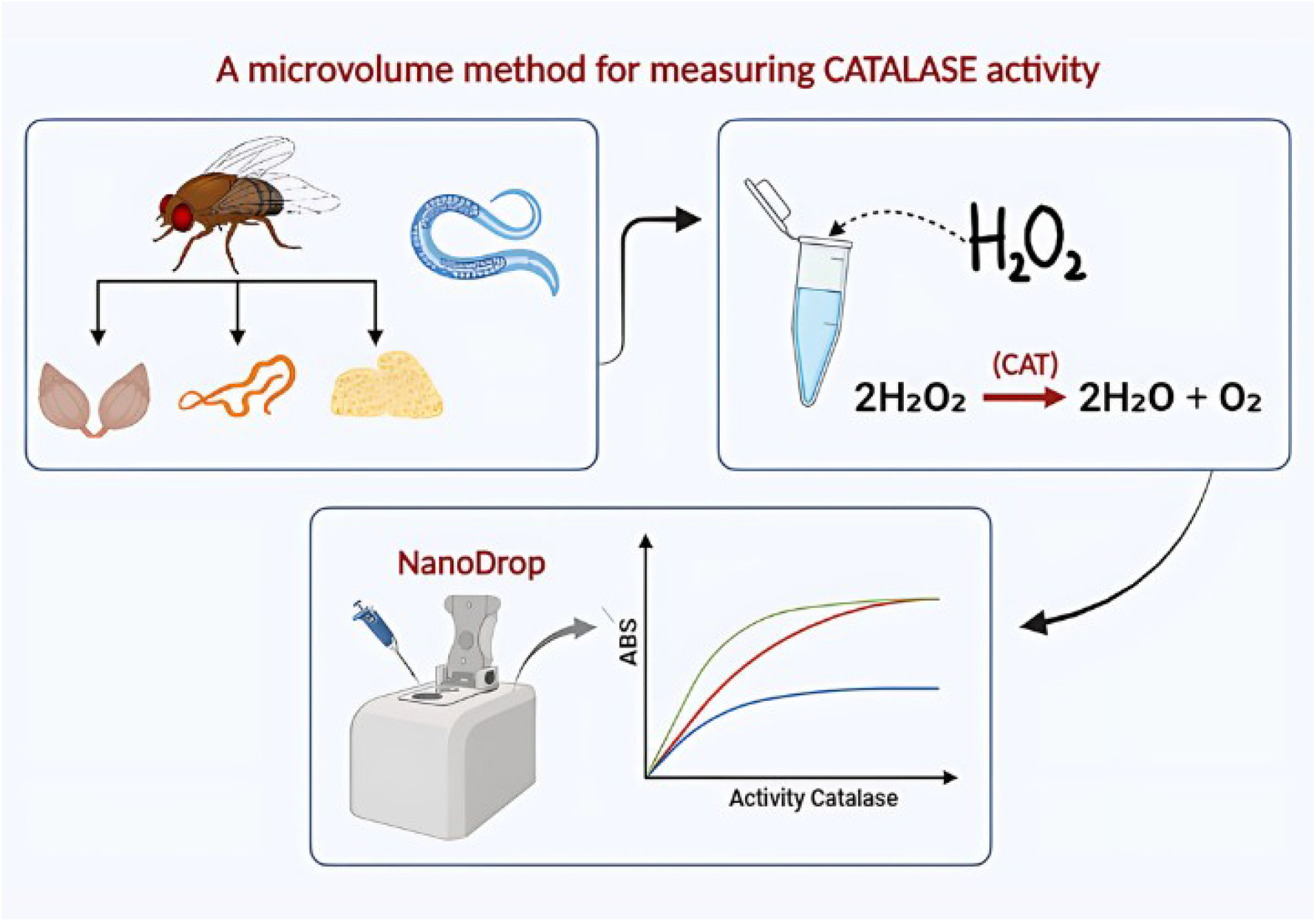

## 1. Introduction

In a cellular environment characterized by the constant dynamics of redox processes, hydrogen peroxide (H_2_O_2_) emerges as a key molecule for the maintenance of oxidative balance (Lennicke and Cocheme 2021; Sies et al. 2024). As an unavoidable by-product of aerobic metabolism, it reflects the intrinsic duality of reactive oxygen species (ROS), which are indispensable as mediators of cellular signaling in moderate concentrations, but are potentially harmful when accumulated to excess (Courgeon et al. 1988; Rabah et al. 2025, Oswald et al 2018).

The toxicity of H_2_O_2_ becomes evident when its production exceeds the antioxidant capacity of the cells (Sies 2020). Under these conditions, the excess of H_2_O_2_ can induce significant oxidative stress, especially through its conversion into highly reactive hydroxyl radicals (•OH), which indiscriminately attack biomolecules (Birben et al. 2012; Halliwell et al. 2021). This process occurs via the Fenton reaction: Fe^2^+ + H_2_O_2_ → Fe^3^+ + OH? + •OH, where ferrous iron (Fe^2^+) donates an electron to H_2_O_2_, generating ferric iron (Fe^3^+), a hydroxide ion (OH?), and the hydroxyl radical (•OH) (Koppenol et al. 2001; Fridovich 1986). Thus, the excessive accumulation of H_2_O_2_, by intensifying oxidative stress, compromises the structural and functional integrity of cells. This can facilitate the development of metabolic dysfunctions and pathologies associated with redox overload (Heo et al. 2020; Olufunmilayo et al. 2023; Saito et al. 2006).

To mitigate the harmful effects of H_2_O_2_, aerobic organisms have an antioxidant system, with catalase (H_2_O_2_: H_2_O_2_-oxidoreductase, EC 1.11.1.6) being one of the most important enzymes in this context (Chelikani et al. 2004; Halliwell and Gutteridge 1990). Catalase is an antioxidant enzyme that primarily has the task of decomposing H_2_O_2_ and converting it extremely quickly and efficiently into water and molecular oxygen, as described by the following equation: 2 H_2_O_2_ → 2 H2O + O2. Its activity is directly related to the maintenance of redox homeostasis and the response to environmental stressors, such as toxic and genotoxic substances and extreme conditions (Calabrese et al. 2009; Willekens et al. 1997).

Given the importance of catalase in various fields of biological research, analyzing its activity in limited samples is a major technical challenge. Conventional protocols such as spectrophotometry, volumetric assays, colorimetric assays, and the Warburg method are widely used and frequently adapted (Sinha 1972; Jing 2017; Hadwan 2018; Goldstein 1968; Warburg et al. 1927). However, these methods usually require larger sample volumes and the use of multiple reagents.

For small organisms such as nematoid worms like *Caenorhabditis elegans* or *Panagrellus redivivus*, which are about 1.0 to 1.5 mm long and 50 μm in diameter (Stefanello and Loreto 2022; Brenner 1974), the analysis of catalase activity faces the additional challenge of extracting sufficient amounts of protein for accurate measurements. Due to the small size of these organisms, the limited amount of protein extracted compromises the reliability of the results and prevents direct quantification of enzymatic activity.

In this context, the protocol validated in the present study uses the NanoDrop 2000® spectrophotometer for the analysis of catalase activity, focusing on microvolume samples (Baginski and Sommerhalter 2017). In contrast to assays performed in microplates, which are commonly used to increase throughput, reduce costs and enable large-scale analyses, our protocol is specifically designed for scenarios where biological material is limited. The main advantage of this approach is that it allows the analysis of very small samples with sample volumes between 1 and 2 μL without compromising the precision and sensitivity of the measurements. In contrast, microplate assays usually require larger sample volumes and often dilute enzyme concentrations. Originally developed for the quantification of nucleic acids and proteins, this device offers high sensitivity and precision and allows measurements in extremely small volumes of 1 to 2 μL (Desjardins and Conklin 2010; Desjardins et al. 2009). Equipped with UV-VIS technology, the NanoDrop ® covers a broad wavelength range from 190 nm to 840 nm and enables the analysis of various biochemical compounds with excellent spectral resolution (Desjardins and Conklin 2010). Although we have validated this protocol with Nanodrop®, we believe that other nanospectrophotometers such as DeNovix® and Implen NanoPhotometer® will provide similar results.

The protocol described here includes two main assays: (1) the observation of the decay of catalytic activity over time to determine the linearity of the reaction (time curve), and (2) the analysis of the relationship between protein concentration and catalytic activity by performing serial dilutions (protein curve to determine the linearity of enzyme activity as a function of protein concentration). The implementation of this method allows the direct measurement of enzymatic activity in specific tissues and organs of model organisms such as *Drosophila melanogaster* and nematodes such as *C. elegans* and *Panagrellus redivivus*, whose characteristics make conventional analysis with traditional spectrophotometric cuvettes difficult. The proposed protocol thus fills a gap in catalase analysis for small biological samples and in volume-limited contexts,

## 2. Materials and methods

### 2.1) Chemicals and reagents

Phosphate buffered saline (PBS) consisted of 37 mM sodium chloride (NaCl), 2.7 mM potassium chloride (KCl), 10 mM disodium hydrogen phosphate (Na_2_HPO_4_) and 1.8 mM potassium dihydrogen phosphate (KH_2_PO_4_), pH 7.5. The salts were purchased from Sigma Aldrich and were of analytical grade. Sodium dodecyl sulfate (SDS) 1% was preceded with Fisher Scientific electrophoresis grade reagent. Hydrogen peroxide (H_2_O_2_) 20 mM solution was prediluted with analytical grade H_2_O_2_ from Exodo Científica. Diethyl ether (CH3CH_2_)_2_O was purchased from Dinâmica Química, analytical grade.

### 2.2) Organisms used and samples preparation

Organs of *Drosophila melanogaster*, strain Oregon R, were used for the assays. The fruit flies were anesthetized with diethyl ether and dissected in a drop of PBS under the stereomicroscope. Adult females were used to test the ovaries and malpighian tubules. Third instar larvae were used for the fat body. The assays were also carried out with the whole body of the worm *Panagrellus redivivus* (Nematoda).

For the sample preparation of *D. melanogaster*, four ovaries, about 60 Malpighian tubules, and the fat body of five larvae were added separately in 100 μL PBS with 1 μL SDS in a 1.5 mL microtube. The material was homogenized with microtube polypropylene pestles. The samples were then centrifuged at 5500 g for 5 minutes. Subsequently, 64 μL of the supernatant was transferred to a new tube, which was used for the following steps. The experiments were carried out in triplicate. A procedure similar to that described for *D. melanogaster* was used for *Panagrellus redivivus*. In this case, ∼400 worms were homogenized for one sample.

### 2.3) Protein quantification

Protein quantification was performed with the NanoDrop at 280 nm according to the manufacturer’s instructions. PBS 1X was used as a blank for all measurements. For each sample, 2 μL of the supernatant was used for the initial protein quantification. For diluted samples, the protein concentration was remeasured after each dilution step to ensure accuracy prior to enzymatic assays.

### 2.4) Catalase Enzyme Assays (CAT)

The enzymatic activity of catalase was measured using the UV-Vis mode of the NanoDrop at 240 nm. The device was calibrated with 2 μL 1X PBS as blank. An initial absorbance measurement was taken before the addition of H_2_O_2_ and later subtracted from the final absorbance values to correct for background interference. The experimental conditions were maintained at 25°C in an air-conditioned room.

To evaluate catalase activity over time, 20 μL of 20 mM H_2_O_2_ was added to 60 μL of homogenate, resulting in a final H_2_O_2_ concentration of 5 mM. The first absorbance measurement was performed immediately after the addition of H_2_O_2_ to establish the reference value for time 0 minutes. Enzyme activity was monitored over a period of 10 minutes, with measurements taken every 1 minute. The decrease in absorbance over time was interpreted as the rate of H_2_O_2_ decomposition. Each sample was tested in triplicate, and three independent measurements were taken per sample to ensure reproducibility and accuracy of the data. A control with PBS and H_2_O_2_, without the addition of protein, was included in all assays.

To evaluate catalase activity at different protein concentrations, the homogenates were diluted in a 1:1 ratio in PBS buffer. This dilution was repeated in five successive steps, resulting in samples with progressively lower protein concentrations. After each dilution, the protein concentration was re-measured using the NanoDrop 2000 to confirm the actual protein levels before proceeding with the enzyme assays. For this assay, 30 μL of homogenate was mixed with 10 μL of 20 mM H_2_O_2_, maintaining a homogenate to H_2_O_2_ ratio of 3:1 and a total reaction volume of 40 μL per sample. The absorbance of 2 μL of each sample was measured exactly 1 minute after the addition of H_2_O_2_. This procedure was repeated with three independent measurements per sample.

In order to compare the enzymatic efficiency between the different tissues, a normalization calculation (or data fitting) was performed, as each tissue differs significantly in the amount of protein required for the tests (specific activity). This allowed the normalization of the activity values and the graphical representation of the reaction curves. This procedure ensured that the differences observed reflected the specific catalase activity and not differences in protein concentration

### 2.5) Inhibitors and interveners

To test whether the measured activity is exclusively due to catalase, assays were performed with azide (NaN_3_), a known catalase inhibitor (Mueller et al. 1997; Montavon, et al. 2007; Iwase et al. 2013), at final concentrations of 0.4 and 2 mM. The ovaries of *Drosophila melanogaster* and the nematode *Panagrellus redivivus* were used. The same amount of tissue used in the enzyme assays was homogenized and divided into two parts: one part was treated with azide, and the other was kept as a control. The same procedure was followed for both organisms.

To investigate the possible influence of SDS on catalase activity (Wang et al, 2016), *P. redivivus* and *D. melanogaster* ovaries were analyzed. One set of *P. redivivus* samples was homogenized with SDS, while another was homogenized with liquid nitrogen (-196°C) after two freeze-thaw cycles to ensure effective homogenization without detergent. The same procedure was applied to *D. melanogaster* ovaries. To assess whether SDS interfered with catalase activity, the samples were compared after adjusting for protein concentration.

## 3. Results

To analyze the sensitivity of the NanoDrop spectrophotometer for hydrogen peroxide quantification, different concentrations of H_2_O_2_ were measured at variable wavelengths (Fig. 1). Above 250 nm, absorbance values rapidly decreased with dilution and became negligible after the fourth dilution. In contrast, 230 nm and 240 nm exhibited high initial absorbance that remained detectable even at lower concentrations, indicating that the peak absorbance of H_2_O_2_ occurs within this range. These findings align with the classical methods which quantifies catalase activity at 240 nm in the presence of phosphate buffer (Aebi 1984).

**Figure 1.**
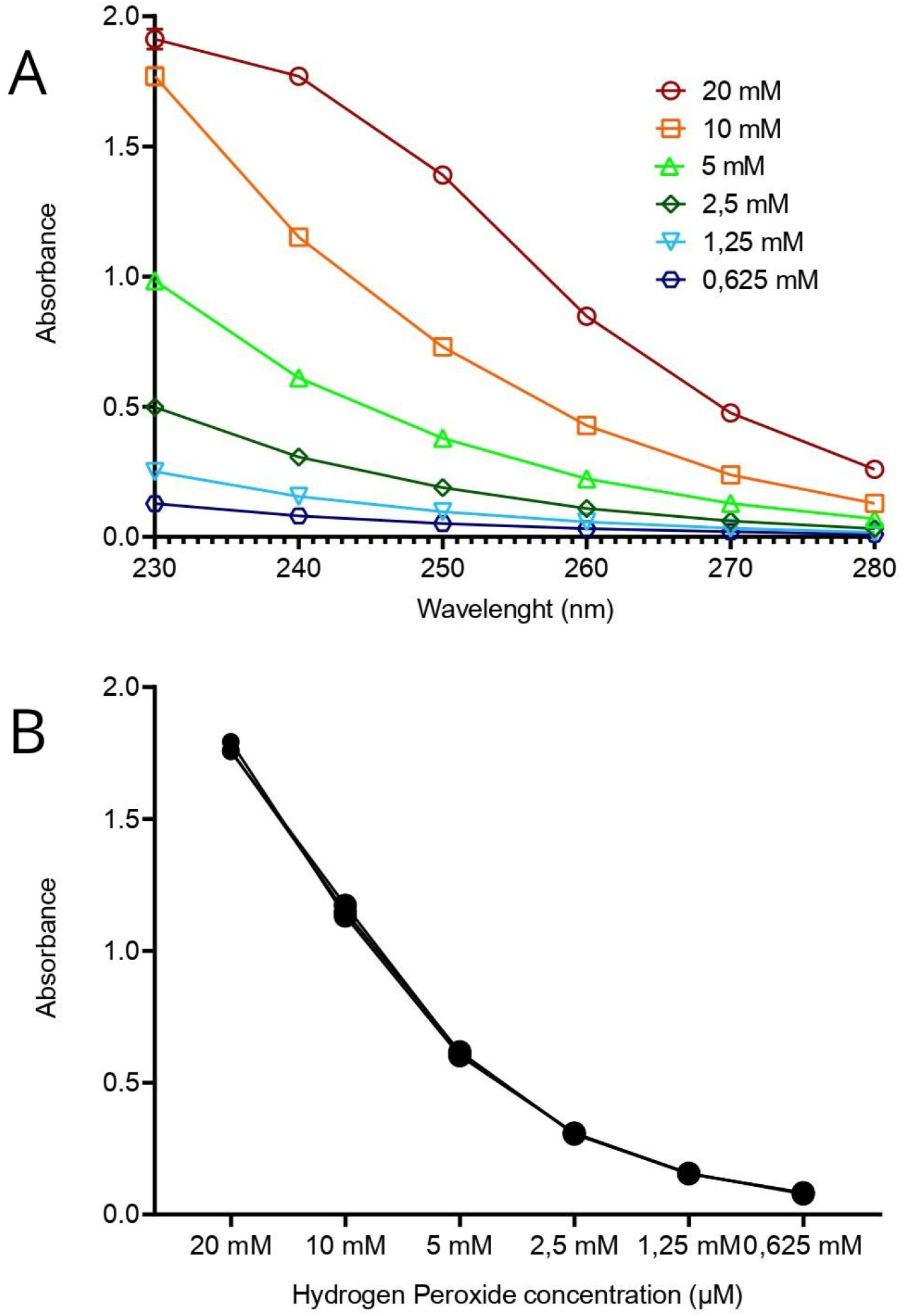
Absorbance of hydrogen peroxide at different wavelengths and concentrations: (A) Absorbance at different concentrations of H_2_O_2_ between 230-280 nm, highlighting the greater stability and sensitivity at 230 nm and 240 nm. (B) Reduction in absorbance at 240 nm with decreasing concentration of H_2_O_2_.

When monitoring hydrogen peroxide degradation over time (Fig. 2A), significant differences in catalase activity were observed among the analyzed tissues. Malpighian tubules showed the highest catalytic activity, with H_2_O_2_ consumption following a linear pattern for approximately 4 to 5 minutes (p < 0.0001). The fat body displayed intermediate catalytic activity, while ovaries required substantially higher protein concentrations to reach detectable activity levels. The control group (PBS + H_2_O_2_) exhibited no significant change in absorbance over 10 minutes (p = 1.000), confirming that the observed changes were catalase-dependent.

**Figure 2.**
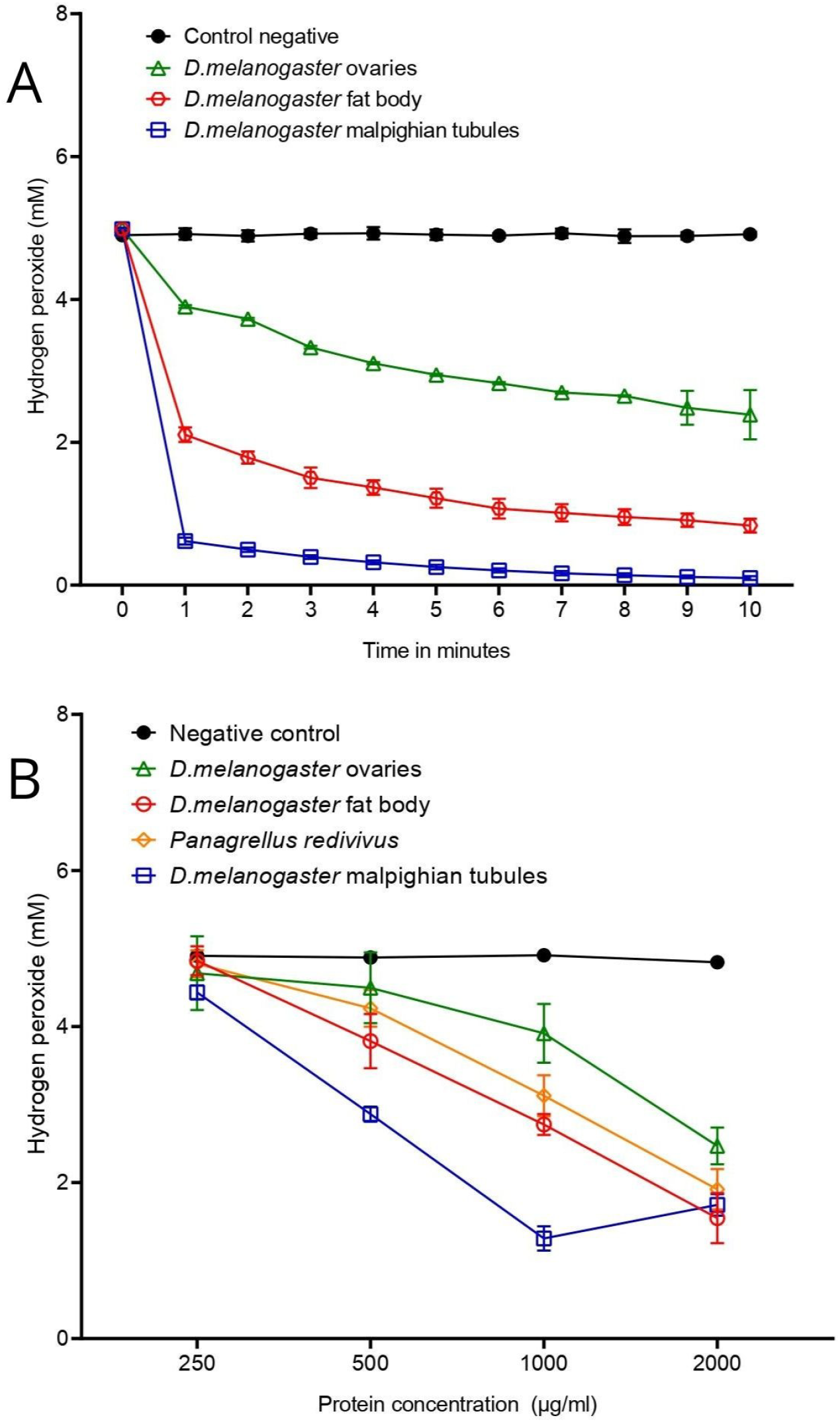
Enzymatic activity of catalase: (A) Decomposition of H_2_O_2_ over time in *D. melanogaster* tissues. Malpighian tubules showed the highest activity, followed by the fat body. (B) Comparison of catalase activity in different tissues at different protein dilutions. Malpighian tubules showed the highest activity, followed by fat bodies and ovaries. *P. redivivus* showed intermediate activity.

Tests in which absorbance was measured for different protein dilutions at a fixation time of one minute confirmed different patterns of catalytic activity in the different tissues (Fig. 2B). Malpighian tubules showed significantly higher activity, especially at higher protein concentrations. From 1000 μg protein/ml, the Malpighian tubules consistently showed higher activity (p < 0.0001). The fat body showed mean activity that was significantly different from the ovaries at all dilutions (p < 0.0001). This suggests fat body activity is robust but lower than Malpighian tubules and higher than ovaries. Ovaries required higher protein concentrations for detectable activity. Comparisons with the negative control (PBS + H_2_O_2_) showed no significant differences at higher dilutions (p > 0.9), indicating minimal or absent activity. Only at higher concentrations (2000 μg protein/ml and 1000 μg protein/ml) did the ovaries differ significantly from the control (p < 0.0001), indicating low catalytic efficiency.

The nematode *Panagrellus redivivus* showed catalytic activity similar to that of the Malpighian tubules and the fat body. Significant differences were observed at all concentrations compared to the ovaries. Starting at a concentration of 1000 μg protein/ml, *Panagrellus* showed a more efficient enzyme. The negative control (PBS + H_2_O_2_) did not show significant absorbance, confirming that the differences in the other groups were due to catalase activity.

Catalase activity was completely inhibited by 0.4 and 2 mM of azide, indicating that the activity was not related to other peroxidases such as glutathione peroxidases and peroxiredoxins. The heat-treated enzyme was included as an additional control. The degradation of hydrogen peroxide depends on the integrity of the enzyme (Fig. 3). The catalase decreased in absorbance at 240 nm as a function of time, indicating the decomposition of hydrogen peroxide. In contrast, azide and heat-denatured catalase did not decompose the peroxide and exhibited absorbance values similar to the negative control (PBS + H_2_O_2_). These differences between active catalase and the other groups were statistically significant (ANOVA, p < 0.0001).

**Figure 3.**
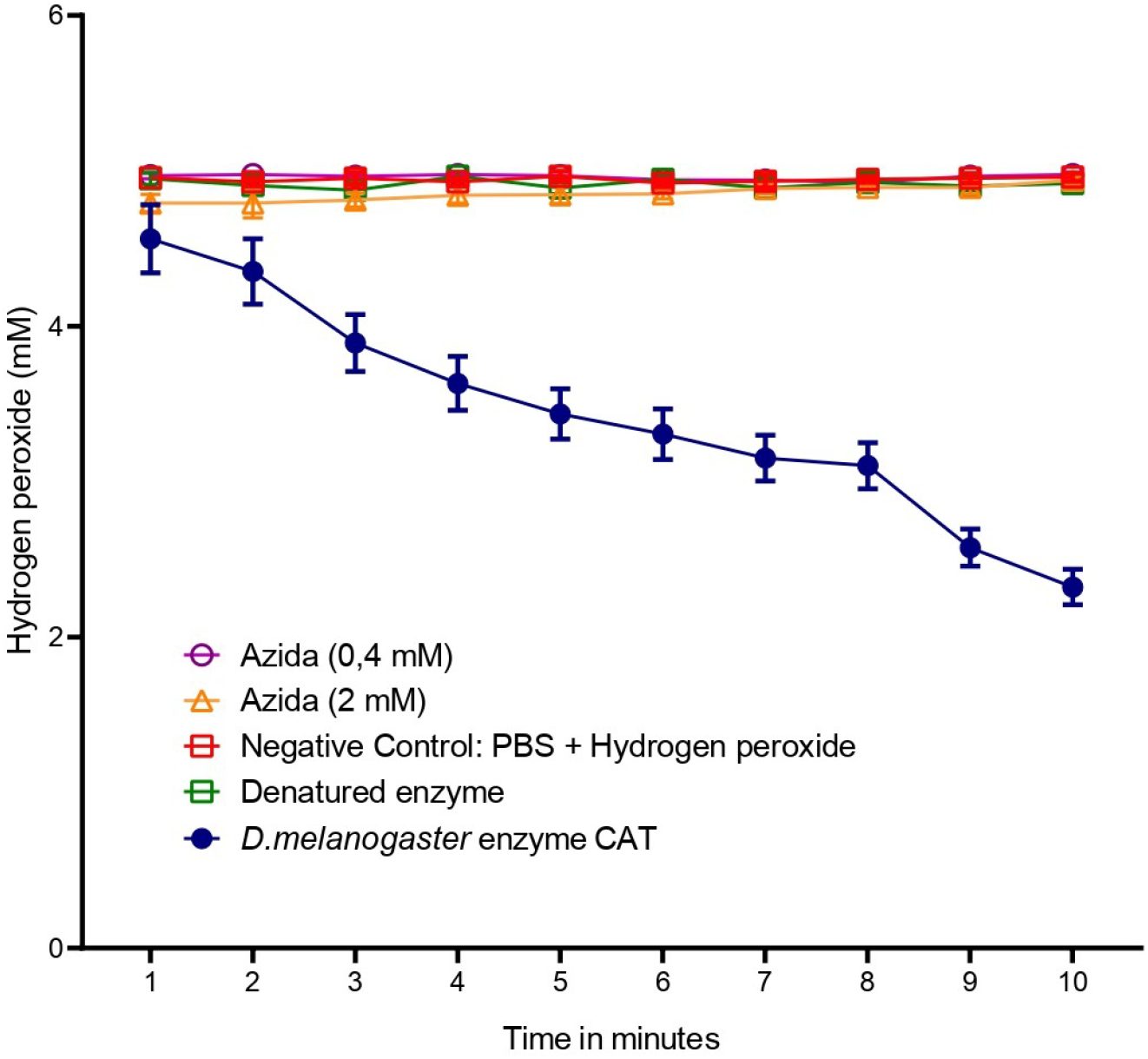
Inhibition of catalase activity: Absorbance measured over time to evaluate the degradation of H_2_O_2_ by active and inhibited catalase. The groups treated with azide (0.4 mM and 2 mM) and denatured catalase showed stable values, similar to the negative control (PBS + H_2_O_2_), demonstrating the absence of catalytic activity.

Tests were performed to analyze whether catalase activity could be modified by the sodium dodecyl sulfate (SDS) used in the homogenizer buffer (Fig. 4). In general, SDS exhibited higher enzymatic activity due to more effective hydrogen peroxide degradation when compared to the activity determined by homogenizing the tissues with freezing-and-thawing the samples in liquid nitrogen. Similar to the results obtained with flies, *P. redivivus*, catalase activity was higher with SDS than after freezing-thawing the worms (p < 0.05), suggesting better enzyme extraction from tissues or by stabilizing or activating catalase, particularly at 1500 e 1000 μg/ml (p < 0.05). Similarly, *D. melanogaster* ovaries exhibited greater enzymatic activity with SDS at the higher protein concentrations tested (p < 0.01). These results suggest that SDS at low concentrations did not inhibit catalase and may have increased activity by improving protein extraction efficiency or by stabilizing the enzyme.

**Figure 4.**
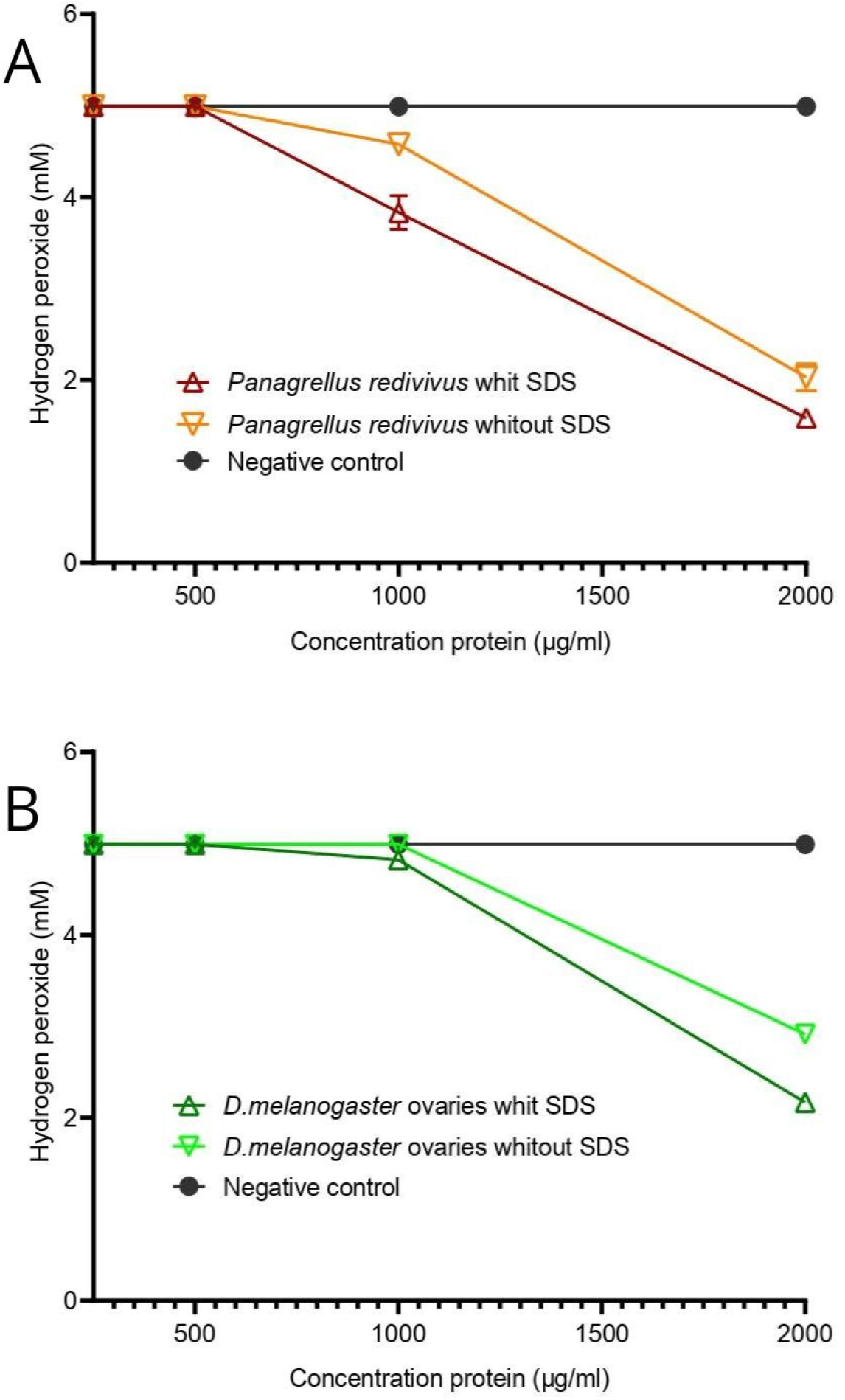
Effect of SDS on catalase activity in *Panagrellus redivivus* and *Drosophila melanogaster* ovaries. The use of SDS resulted in higher enzymatic activity compared to homogenization by freezing and thawing. This effect was more pronounced at protein concentrations of 1500 and 1000 μg/ml, suggesting that SDS may improve the efficiency of extraction or stabilization of catalase.

## 4. Discussion

UV-VIS spectrophotometry is widely used to monitor the activity of enzymes such as catalase (CAT) due to its high sensitivity in measuring variations in H_2_O_2_ concentration. Some of the protocols already described suggest adaptation for reduced volumes, which is particularly useful when large quantities of biological samples are available and allows for a more practical and cost-effective procedure (Hadwan et al. 2024; Kadhum and Hadwan 2021). While this approach is advantageous in situations with high sample availability. The protocol proposed here excels in scenarios where the amount of biological material is limited, such as studies with small organisms and reduced tissue samples, as it is able to detect enzymatic activity with only 2 μL of sample.

The results of this study show that the NanoDrop spectrophotometer has greater stability and sensitivity at wavelengths of 230 nm and 240 nm, ideal for monitoring H_2_O_2_ decay. Previously, Baginski and Sommerhalter (2017) measured catalase activity using the NanoDrop® spectrophotometer. However, they used established protocols for conventional spectrophotometers. Here, we developed and validated an innovative protocol using microvolumes and the NanoDrop spectrophotometer. Other nanospectrophotometers such as the DeNovix® and the Implen NanoPhotometer® are likely to provide similar results.

The values obtained for higher sensitivity wavelengths are consistent with the literature, which highlights the absorption peak of H_2_O_2_ at 230 nm, while 240 nm isnm is considered optimal for catalase, as it ensures good H_2_O_2_ at absorption while minimizing interference from other compounds, such as reaction products (Beers and Sizer 1952). Above 240 nm, a significant decrease in absorbance was observed with the dilution of the sample, which reduces the sensitivity and reliability of the measurements, as Beers and Sizer (1952) found.

An important adjustment was the initial concentration of H_2_O_2_, which was set to 5 mM based on the adjustment proposed by Baginski and Sommerhalter (2017) for the use of the NanoDrop. In contrast to conventional spectrophotometers, which use cuvettes with a fixed optical path length of 10 mm, the NanoDrop measures samples with a much smaller optical path length, typically between 0.5 and 1 mm. This difference can affect the absorbance intensity as the absorbance is proportional to the optical path length. Therefore, a shorter path can result in lower readings for the same H_2_O_2_ concentration (Johnson and Goody 2011). To keep the initial absorbance within the detection range of the device and to ensure precise monitoring of the reaction kinetics, we adjusted the H_2_O_2_ concentration accordingly.

We have observed that CAT efficiency is reflected in the lower amount of protein required to achieve measurable activity in tissues with high catalytic efficiency, such as the Malpighian tubules. This phenomenon can be explained by the principles of enzyme kinetics, in particular the Michaelis–Menten equation, which relates the reaction rate to the concentrations of substrate and enzyme (Johnson and Goody 2011). Therefore, the specific activity is higher in more efficient tissues, leading to robust results even with reduced biological material. This relationship emphasises the importance of adjusting the amount of protein to ensure the validity and reproducibility of the assay.

In the catalase enzyme activity tests, the Malpighian tubules showed the highest enzymatic efficiency, followed by the fat bodies, which showed intermediate activity, and the ovaries, which showed the lowest activity. These results are consistent with the physiology of each tissue and support studies indicating the role of these tissues in osmotic homeostasis, excretion of metabolites and detoxification processes, mainly in the Malpighian tubules (Dow 2009; Giglio and Brandmayr 2017; Stergiopoulos et al. 2009; Li et al, 2018). The detection of CAT activity in *Panagrellus redivivus* shows that the methodology can be applied to worms similar in size to *Caenorhabditis elegan*s and other small organisms.

The use of sodium azide can aid in assessing the specificity of catalase (CAT), as no H_2_O_2_ degradation was observed in the treated groups, suggesting that the activity detected was mainly from this enzyme. This result is consistent with Keilin and Hartree (Keilin and Hartree 1945), who described the formation of a stable complex between azide and CAT without significantly altering its structure. However, azide is not completely selective, as it can inhibit other heme-dependent enzymes and generate reactive azidyl radicals that may interfere with assays (Kalyanaraman et al, 1985). To distinguish catalase activity from possible interference, we performed an additional experiment with a heat-denatured enzyme. The complete loss of catalytic activity indicates that H_2_O_2_ degradation is directly linked to the structural integrity of CAT. Studies suggest that heat affects the functionality of enzymes and serves as a complementary method to confirm their specific activity (Hook and Harding 2004; Eyster 1950). The comparison of chemical inhibition by azide with thermal denaturation therefore supports the hypothesis that the observed effects on H_2_O_2_ degradation are predominantly due to catalase activity and minimize the influence of secondary enzymes.

Although SDS is a surfactant with potentially denaturing properties, low concentrations did not affect the catalytic activity of CAT. Wang et al. (2016) indicate that SDS concentrations between 0.1 μM and 0.2 mM do not significantly affect CAT activity in mouse hepatocytes. In the present study, the final SDS concentration was approximately 0.0143%, a value below the threshold associated with CAT inhibition (≥ 0.5 mM), which acts as a solubilizer without affecting enzymatic activity.

## 5. Conclusion

This study highlights the effectiveness of the NanoDrop spectrophotometer in quantifying catalase activity in microvolumes, demonstrating that it is a simple and cost-effective technique. The methodology, which allows both protein measurement and enzyme activity assessment to be performed using a single instrument, proved advantageous for research involving small organisms or even different parts of organisms such as *Drosophila melanogaster* or small worms as *Panagrellus redivivus*. We detected variations in catalase activity among the tissues analyzed and organisms analyzed. Thus, this approach represents a practical and precise alternative for investigating oxidative stress and redox homeostasis in small organisms, organs, and tissues.

## 6. Credit authorship contribution statement

**Beatriz Corrêa da Rocha:** investigation, methodology, formal analysis, writing–original draft, data curation. **Marcos Trindade Da Rosa**, investigation, formal analysis, writing– original draft, data curation; **João Batista Teixeira Da Rocha:** formal analysis, writing– original draft, **Elgion Lucio Silva Loreto:** conceptualization, writing–original draft, supervision, writing–review and editing. All authors have made intellectual contributions to the research project and approved the final manuscript.

## 7. Declarations

### Ethics approval

This study did not involve human or chordate animal experimentation and therefore did not require ethics committee approval under local law.

### Consent to participate

Not applicable.

### Consent to publish

Not applicable

## 9. Data availability

Data supporting the findings reported in this study are available upon reasonable request.

## Bibliography

Aebi H (1984) Catalase in vitro, Methods Enzymol. 105:121–126.

Baginski R, Sommerhalter M (2017) A manganese catalase from Thermomicrobium roseum with peroxidase and catecholase activity, Extremophiles 21: 201–210, 10.1007/s00792-016-0896-9.

Beers Jr. RF, Sizer IW (1952) A spectrophotometric method for measuring the breakdown of hydrogen peroxide by catalase, J. Biol. Chem. 195:133–140, 10.1016/S0021-9258(18)53640-0.

Birben E, Sahiner UM, Sackesen C, Erzurum S, Kalayci O. (2012) Oxidative stress and antioxidant defense, World Allergy Organ. J. 5: 9–19, 10.1097/WOX.0b013e3182439613.

Brenner S (1974) The genetics of Caenorhabditis elegans, Genetics 77: 71–94.

Calabrese V, Cornelius C, Mancuso C, Lentile R, Stella AMG, Butterfield DA (2009) Redox homeostasis and cellular stress response in aging and neurodegeneration, in: S.A. Mansouri (Ed.), Free Radicals and Antioxidant Protocols, 2nd ed., New York: Springer, 2009, pp. 285–308, 10.1007/978-1-60327-029-8_17.

Chelikani P, Fita I, Loewen PC (2004) Diversity of structures and properties among catalases, Cell. Mol. Life Sci. 61:192–208, 10.1007/s00018-003-3206-5.

Courgeon AM,]Rollet E, Becker J, Maisonhaute C, Best-Belpomme M (1988) Hydrogen peroxide (H_2_O_2_) induces actin and some heat-shock proteins in Drosophila cells, Eur. J. Biochem. 171: 163–170, 10.1111/j.1432-1033.1988.tb13772.x.

Desjardins P, Conklin D (2010) NanoDrop microvolume quantitation of nucleic acids, J. Vis. Exp. 45:e2565.

Desjardins P, Hansen JB, Allen M (2009) Microvolume protein concentration determination using the NanoDrop 2000c spectrophotometer, J. Vis. Exp. 33: e1610, 10.3791/1610.

Dow JAT (2009) Insights into the Malpighian tubule from functional genomics, J. Exp. Biol. 212: 435–445, 10.1242/jeb.024224.

Eyster HC (1950) Effect of temperature on catalase activity, Ohio J. Sci. 50: 273–277.

Fridovich I (1986) Biological effects of the superoxide radical, Arch. Biochem. Biophys. 247: 1–11, 10.1016/0003-9861(86)90046-3.

Giglio A, Brandmayr P (2017) Structural and functional alterations in Malpighian tubules as biomarkers of environmental pollution: synopsis and prospective, J. Appl. Toxicol. 37:889–894, 10.1002/jat.3454.

Goldstein DB (1968) A method for the assay of catalase with the oxygen cathode, Anal. Biochem. 28:431–437.

Hadwan MH (2018) Simple spectrophotometric assay for measuring catalase activity in biological tissues, BMC Biochem. 19:7, 10.1186/s12858-018-0097-5.

Hadwan MH, Hussein MJ, Mohammed RM, Hadwan AM, Saad Al-Kawaz H, Al-Obaidy SSM, Al Talebi ZA (2024) An improved method for measuring catalase activity in biological samples. Biol Methods Protoc. 9(1):bpae015: doi: 10.1093/biomethods/bpae015.

Halliwell B, Adhikary A, Dingfelder M, Dizdaroğlu M (2021) Hydroxyl radical is a significant player in oxidative DNA damage in vivo, Chem. Soc. Rev. 50: 8355–8360, 10.1039/d1cs00044f.

Halliwell B, Gutteridge JMC (1990) Role of free radicals and catalytic metal ions in human disease: An overview, Methods Enzymol. 186: 1–85, 10.1016/0076-6879(90)86093-b.

Heo S, Kim S, Kang D (2020) The role of hydrogen peroxide and peroxiredoxins throughout the cell cycle, Antioxidants (Basel) 9: 280, 10.3390/antiox9040280.

Hook DWA, Harding JJ (2004) Molecular chaperones protect catalase against thermal stress, Eur. J. Biochem. 247: 380–385.

Iwase T, Tajima A, Sugimoto S, Okuda K, Hironaka I, Kamata Y, Takada K, Mizunoe Y (2013) A simple assay for measuring catalase activity: A visual approach, Sci. Rep. 3: 3081 10.1038/srep03081

Jing D, Wilkinson L, Liu Y (2017) Superoxide dismutase (SOD) and catalase (CAT) activity assay protocols for Caenorhabditis elegans, Bio-protocol 7:e2505, 10.21769/BioProtoc.2505.

Johnson KA, Goody RS (2011) The Original Michaelis Constant: Translation of the 1913 Michaelis–Menten Paper, Biochemistry 50: 8264–8269, 10.1021/bi201284u.

Kadhum MA, Hadwan MH (2021) A precise and simple method for measuring catalase activity in biological samples. Chem. Pap. 75:1669–1678. 10.1007/s11696-020-01401-0

Kalyanaraman B, Janzen EG, Mason RP (1985) Spin trapping of the azidyl radical in azide/catalase/H_2_O_2_ and various azide/peroxidase/H_2_O_2_ peroxidizing systems, J. Biol. Chem. 260: 4003–4006.

Keilin D, Hartree EF (1945) Properties of azide-catalase, Biochem. J. 39:148–156.

Koppenol WH (2001) The Haber-Weiss cycle – 70 years later, Redox Rep. 6: 229–234, 10.1179/135100001101536373.

Lennicke C, Cocheme HM (2021) Redox metabolism: ROS as specific molecular regulators of cell signaling and function, Mol. Cell 81:3977–3992, 10.1016/j.molcel.2021.08.018.

Li W, Young JF, Sun J (2018) NADPH oxidase-generated reactive oxygen species in mature follicles are essential for Drosophila ovulation, Proc. Natl. Acad. Sci. (PNAS) 115:7765–7770, 10.1073/pnas.1800115115.

Montavon P, Kukic KR, Bortlik K (2007) A simple method to measure effective catalase activities: Optimization, validation, and application in green coffee, Anal. Biochem. 360: 207–215.

Mueller S, Riedel HD, Stremmel W (1997) Determination of catalase activity at physiological hydrogen peroxide concentrations, Anal. Biochem. 245: 55–60.

Olufunmilayo EO, Gerke-Duncan MB, Holsinger RMD (2023) Oxidative stress and antioxidants in neurodegenerative disorders, Antioxidants 12: 517, 10.3390/antiox12020517.

Oswald MCW, Brooks PS, Zwart MF, Mukherjee A, West RJH, Giachello CNG, Morarach K, Baines RA, Sweeney ST, Landgraf M (2018) Reactive oxygen species regulate activity-dependent neuronal plasticity in Drosophila, eLife 7: e39393, 10.7554/eLife.39393.

Rabah Y, Berwick JP, Sagar N, et al. (2025) Astrocyte-to-neuron H_2_O_2_ signalling supports long-term memory formation in Drosophila and is impaired in an Alzheimer’s disease model, Nat. Metab. 7:321–335, 10.1038/s42255-024-01189-3.

Saito Y, Liang G, Egger G, Friedman JM, Chuang JC, Coetzee GA, Jones PA (2006) Specific activation of microRNA-127 with downregulation of the proto-oncogene BCL6 by chromatin-modifying drugs in human cancer cells, Cancer Cell 9: 435–443, 10.1016/j.ccr.2006.04.020.

Sies H, Mailloux RJ, Jakob U (2024) Fundamentals of redox regulation in biology, Nat. Rev. Mol. Cell Biol. 25 701–719, 10.1038/s41580-024-00730-2.

Sies S (2020) Hydrogen peroxide as a central redox signaling molecule in physiological oxidative stress: Oxidative eustress, Free Radic. Biol. Med. 160:47–58, 10.1016/j.freeradbiomed.2020.09.021.

Sinha AK (1972) Colorimetric assay of catalase, Anal. Biochem. 47: 389–394, 10.1016/0003-2697(72)90132-7.

Stefanello L, Loreto ELS (2022) Panagrellus redivivus: a promising nematode for cell biology practical classes, Research, Society and Development 11: e29311225629, 10.33448/rsd-v11i2.25629.

Stergiopoulos K, Cabrero P, Davies SA, Dow JA (2009) Salty dog, an SLC5 symporter, modulates Drosophila response to salt stress, Physiol. Genomics 37:1–11, 10.1152/physiolgenomics.90360.2008.

Wang J, Wang J, Xu C, Liu R, Chen Y (2016) Molecular mechanism of catalase activity change under sodium dodecyl sulfate-induced oxidative stress in the mouse primary hepatocytes, J. Hazard. Mater. 307:173–183, 10.1016/j.jhazmat.2015.11.060.

Warburg O, Wind F, Negelein E (1927) The metabolism of tumors in the body, J. Gen. Physiol. 8 519–530, 10.1085/jgp.8.6.519.

Willekens H, Chamnongpol S, Davey M, Schraudner M, Langebartels C, Van Montagu M, Inzé D, Van Camp W (1997) Catalase is a sink for H_2_O_2_ and is indispensable for stress defence in C_3_ plants, EMBO J. 16:4806–4816, 10.1093/emboj/16.16.4806.

